# Investigating the accuracy of imputed genotypes in Nellore cattle using the ARS-UCD1.2 assembly of the bovine genome

**DOI:** 10.1101/2020.06.26.172353

**Authors:** Isis da Costa Hermisdorff, Raphael Bermal Costa, Lucia Galvão de Albuquerque, Hubert Pausch, Naveen Kumar Kadri

## Abstract

**Background:** Imputation accuracy among other things depends on the size of the reference panel, the marker’s minor allele frequency (MAF), and the correct placement of variants on the reference genome assembly. Using high-density genotypes of 3938 Nellore cattle from Brazil, we investigated the accuracy of imputation from 50K to 777K SNP density, using map positions determined according to the bovine genome assemblies UMD3.1 and ARS-UCD1.2. We assessed the effect of reference and target panel sizes on the pre-phasing-based imputation quality using ten-fold cross-validation. Further, we compared the reliability of the model-based imputation quality score (R*sq*) from Minimac3 to empirical imputation accuracy.

**Results:** The overall accuracy of imputation measured as the squared correlation between true and imputed allele dosages (R^2^*dose*) was virtually identical using either the UMD3.1 or ARS-UCD1.2 genome assembly. When the size of the reference panel increased from 250 to 2000, R^2^*dose* increased from 0.845 to 0.917, and the number of polymorphic markers in the imputed data set increased from 586,701 to 618,660. Advantages in both accuracy and marker density were also observed when larger target panels were imputed, likely resulting from more accurate haplotype inference. Imputation accuracy and the marker density in the imputed data increased from 0.903 to 0.913 and from 593,239 to 595,570 when haplotypes were inferred in 500 and 2900 target animals, respectively. The model-based imputation quality scores from Minimac3 (R*sq*) were highly correlated to but systematically higher than empirically estimated accuracies. The correlation between these metrics increased with the size of the reference panel and MAF of imputed variants.

**Conclusions:** Accurate imputation of BovineHD BeadChip markers is possible in Nellore cattle using the new bovine reference genome assembly ARS-UCD1.2. The use of large reference and target panels improves the accuracy of the imputed genotypes and provides genotypes for more markers segregating at low frequency for downstream genomic analyses. The model-based imputation quality score from Minimac3 (R*sq*) can be used to detect poorly imputed variants but its reliability depends on the size of the reference panel used and MAF of the imputed variants.

## Background

Genotype imputation is a cost-effective statistical approach to infer genotypes at untyped loci (1–4). Target panels genotyped with low-density SNP chips can be imputed to higher density using a reference panel typed at higher density. Imputation is important to increase the power and resolution of genome-wide association studies (2,5) and the accuracy of genomic breeding values for individuals typed using cheap low-density SNPs genotyping arrays (6). Imputation accuracy depends on several factors including the size and composition of the reference panel (3,7), the minor allele frequency (MAF) of imputed variants (8), and correct placement of variants on the reference genome assembly (7,9,10).

Large and informative reference panels are required to facilitate accurate imputation of rare and low-frequency variants (3,7,11). The effect of the target panel size, on the other hand, has been barely studied. The softwares that implement a pre-phasing-based strategy (12), for example, Minimac3 (13) and IMPUTE2 (14), impute genotypes into phased haplotypes. These tools are probably more sensitive to the size of the target panel because the phasing accuracy depends on the number and the relatedness of the samples considered (15-17). Erroneously inferred haplotypes might compromise the quality of subsequent imputation.

The correct physical placement of the SNPs along the genome is also crucial for accurate phasing and genotype imputation (7,10,18,19). Several iterations of improvement resulted in a highly contiguous and accurate bovine reference sequence (20). However, the correct placement of markers and haplotypes inference remains challenging in repetitive and duplicated regions (21). Recently a highly contiguous version (ARS-UCD1.2) of the bovine reference sequence -- assembled using long sequencing reads -- filled gaps and resolved repetitive regions of the previous UMD3.1 assembly (21).

Identifying and filtering out poorly imputed variants is important to avoid bias in downstream genomic analyses (2). The empirical accuracy of imputation can be obtained by masking and subsequently predicting genotypes in a cross-validation setting. To avoid this computationally demanding and time-consuming process, the per-locus imputation quality scores from imputation programs are often used as indicators for imputation quality. These quality scores are correlated with the true empirical estimates (22), however, their reliability with respect to the size of the reference panel and MAF of the imputed variants has not been tested extensively.

We herein study the quality of imputation from 50K to 777K SNP density in Nellore cattle from Brazil using a pre-phasing-based workflow, implemented using the software packages Eagle2.4 for phasing (16), and Minimac3 for imputation (13). First, we investigate if the improvements in the new bovine genome assembly (ARS-UCD1.2) affect the imputation quality. Next, we test the effect of reference and target panel size on imputation quality. In addition to the commonly used imputation quality metrics, we study the marker density in the imputed data set, i.e. the number of markers segregating in the target panel, after imputation. Finally, we study the reliability of model-based quality scores obtained from the imputation software by comparing it to empirical measures obtained using cross-validation.

## Results

### Comparison of the accuracy of imputation between the UMD3.1 and ARS-UCD1.2 genome assemblies

We considered 3938 Nellore cattle genotyped at 777,962 SNPs using the Illumina BovineHD BeadChip to study the effect of reference genome assemblies on the imputation accuracy. Following quality control (QC) on the genotype data, we determined the position of 684,561 and 683,590 autosomal markers of the Illumina BovineHD BeadChip according to the UMD3.1 and ARS-UCD1.2 bovine genome assemblies, respectively. An intersection of 683,504 autosomal SNPs mapped to both assemblies. The genotypes of 1938 randomly selected animals (target) were reduced to the markers of the Illumina BovineSNP50 genotyping array to mimic low SNP density. The masked genotypes were subsequently inferred *in silico* using 777K genotypes of 2000 animals (reference) in ten-fold cross-validation. The empirical accuracy of imputation was assessed by the squared correlation between true and imputed alleles dosage (R^2^*dose*). There was virtually no difference in imputation accuracy when the markers were aligned to either UMD3.1 or ARS-UCD1.2 with the respective mean R^2^*dose* of 0.916 and 0.917. Only a few SNPs showed large differences in the accuracy of imputation between the two assemblies (Fig. 1a, Additional file 1b). Genotype imputation was less accurate for low-frequency variants (MAF < 5%) (Additional file 1a) using either the ARS-UCD1.2 (0.818) or UMD3.1 (0.817) assembly.

**Figure 1.**
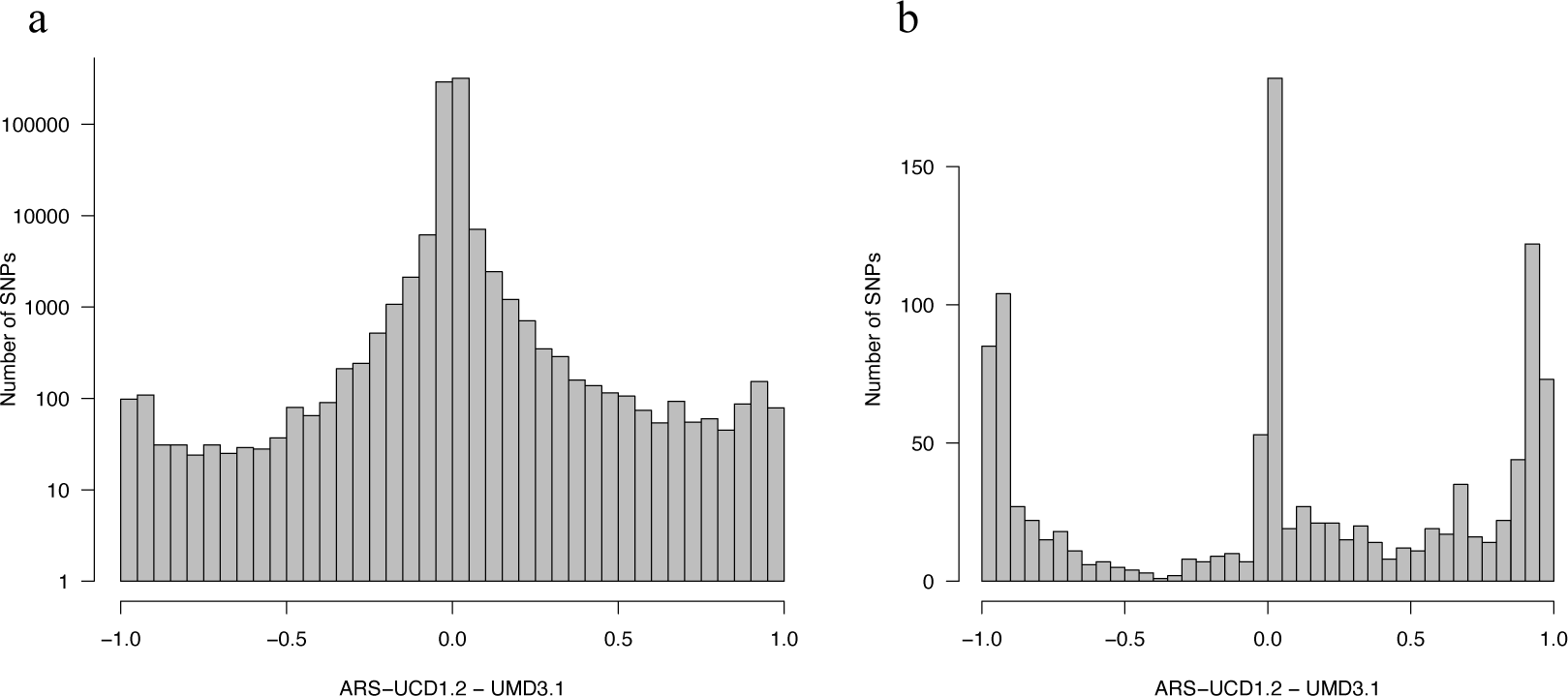
Difference in imputation accuracy according to reference genome assemblies. Difference in imputation accuracy (R^*2*^*dose*) between the ARS-UCD1.2 and UMD3.1 assembly for all makers (a), and for markers that were remapped to a different chromosome (b), when genotypes were aligned to the two Bovine assemblies. To facilitate the interpretation, the y-axis is presented on log-scale in (a).

The new assembly placed 1507 of 1735 markers that were previously unmapped (i.e., not assigned to a chromosome on UMD3.1) to autosomes; 762 of these passed our QC para^−^ meters and segregated in the target panel after imputation. 356 did not pass QC, mainly due to MAF filtering applied, and 389 markers were monomorphic after the imputation. The newly placed autosomal markers were imputed with a mean R^*2*^*dose* value of 0.802. Compared to the UMD3.1 assembly, 2874 markers were placed on a different chromosome in the ARS-UCD1.2 assembly. For 1116 of these markers that passed QC in our data, the mean R^*2*^*dose* was low but improved from 0.515 (UMD3.1) to 0.586 (ARS-UCD1.2) (Fig. 1b).

The analysis of imputation accuracy enabled us to identify regions with poor imputation quality most likely due to misplaced SNPs. We partitioned autosomes into segments of 100 kb and counted the number of SNPs that were poorly imputed despite moderate minor allele frequency (MAF > 1% and R^*2*^*dose* < 0.8). All SNPs in segments with more than three poorly imputed SNPs were considered as putatively misplaced (Fig. 2). A total of 246 segments containing 5447 SNPs (Additional file 2) and 182 segments containing 3549 SNPs (Additional file 3) were identified as misplaced on the UMD3.1 and ARS-UCD1.2 assembly, respectively. A subset of 2205 SNPs was identified as misplaced on both the assemblies with respective mean R^*2*^*dose* of 0.65 and 0.67 when aligned to the UMD3.1 and ARS-UCD1.2 assembly. 3242 SNPs in segments identified as misplaced on the UMD3.1 assembly were imputed with better accuracies (not on poorly imputed segments) on the ARS-UCD1.2 assembly. The mean R^*2*^*dose* for these SNPs increased from 0.78 to 0.93 when aligned to ARS-UCD1.2. The increase in R^*2*^*dose* was even more substantial (from 0.49 to 0.95) for 386 SNPs that were mapped to a different chromosome on the new build. On the other hand, 1344 SNPs -- not on poorly imputed segments when mapped to the UMD3.1 assembly -- were identified as putatively misplaced on the ARS-UCD1.2 assembly. The mean R^*2*^*dose* for these SNPs decreased from 0.89 to 0.73 when mapped to the ARS-UCD1.2 assembly. The drop in the accuracy was even more substantial (from 0.89 to 0.14) for 257 SNPs that were mapped to a different chromosome on the new build.

**Figure 2.**
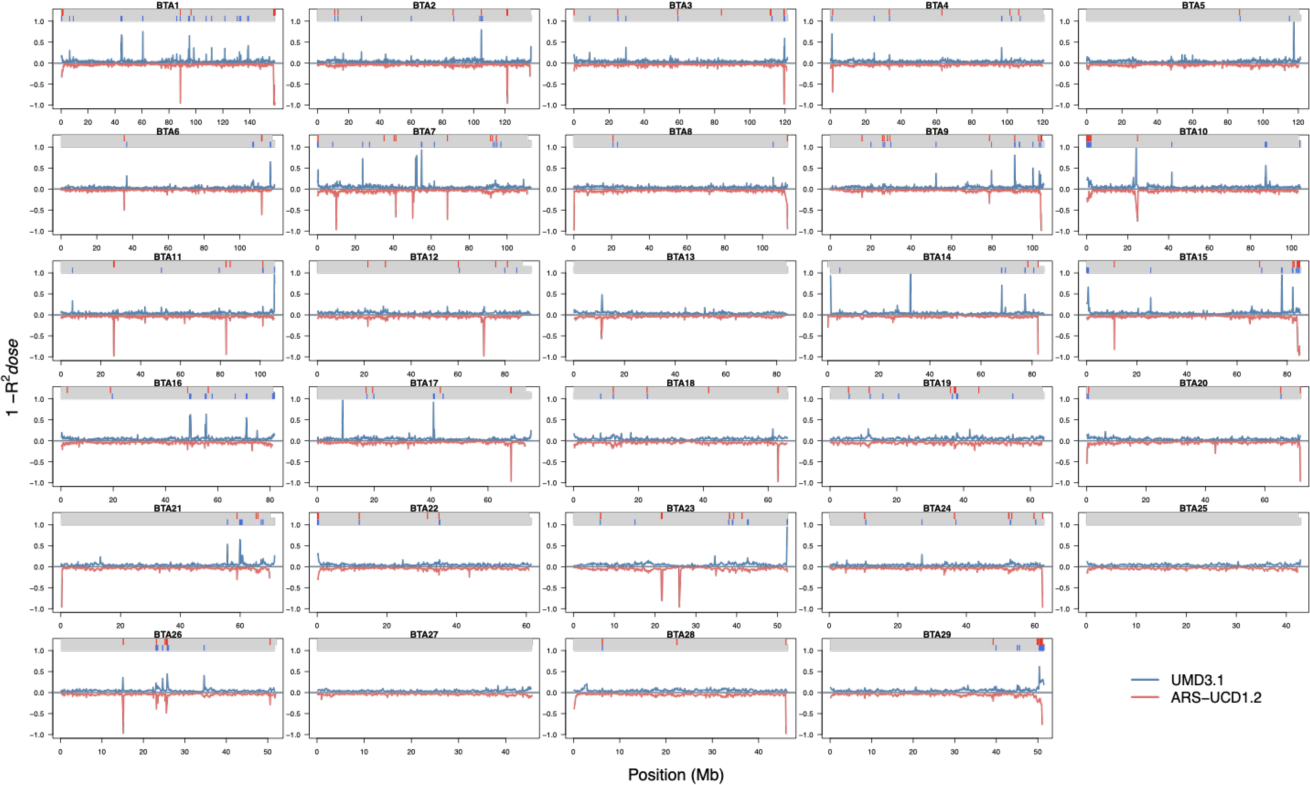
Local imputation accuracy (R*2dose*) when imputing HD genotypes using the two bovine genome assemblies. Mean imputation accuracy (R^*2*^*dose*) for SNPs grouped in segments of 100kb on the 29 autosomes when aligned to the ARS-UCD1.2 and UMD3.1 assemblies. For better visualisation, accuracies are plotted as means of 1-R^*2*^*dose* and R^*2*^*dose*-1 for UMD3.1 and ARS-UCD1.2 assemblies respectively. Number of poorly imputed SNPs (MAF >1% and R*2dose* <0.8) are counted in segments of 100kb. Segments with more than three poorly imputed SNPs are identified as putatively misplaced. The putatively misplaced segments on UMD3.1 (blue) and ARS-UCD3.1 (red) are marked in colored tracks.

### Effect of reference panel size on imputation accuracy and marker density

We used SNPs mapped to ARS-UCD1.2 to study the effect of reference panel size on the accuracy of imputation and marker density in the imputed data set. Using four reference panels containing 250, 500, 1000, and 2000 randomly selected animals, we imputed genotypes for 1938 animals from low (50K) to high density (777K) and compared their imputed and true genotypes using ten-fold cross validation. The mean squared correlation between true and imputed allele dosage (R^*2*^*dose*) increased with an increasing size of reference panels. For instance, R^*2*^*dose* increased from 0.845 to 0.917 (8.5%) when the reference panel size increased from 250 to 2000 (Table 1).

**Table 1.** Empirical accuracies (R^*2*^*dose*, R^*2*^*gt*, CR) and model-based (R*sq*) imputation quality score from Minimac3 using different reference panels

|  |  | Size of the reference panel |  |  |  |
| --- | --- | --- | --- | --- | --- |
|  |  | 250 | 500 | 1000 | 2000 |
| **$R_{sq}$** | **Low-frequency** (MAF < 5%) | 0.829 | 0.856 | 0.871 | 0.876 |
|  | **Common** (MAF > 5%) | 0.942 | 0.960 | 0.971 | 0.978 |
|  | **Global** | 0.909 | 0.928 | 0.938 | 0.944 |
| **$R^2_{dose}$** | **Low-frequency** (MAF < 5%) | 0.688 | 0.737 | 0.778 | 0.818 |
|  | **Common** (MAF > 5%) | 0.909 | 0.936 | 0.953 | 0.965 |
|  | **Global** | 0.845 | 0.874 | 0.897 | 0.917 |
| **$R^2_{gt}$** | **Low-frequency** (MAF < 5%) | 0.690 | 0.746 | 0.792 | 0.832 |
|  | **Common** (MAF > 5%) | 0.887 | 0.920 | 0.941 | 0.956 |
|  | **Global** | 0.832 | 0.868 | 0.895 | 0.917 |
| **CR** | **Low-frequency** (MAF < 5%) | 0.992 | 0.995 | 0.996 | 0.997 |
|  | **Common** (MAF > 5%) | 0.962 | 0.973 | 0.98 | 0.985 |
|  | **Global** | 0.971 | 0.980 | 0.986 | 0.989 |

The increase in R^*2*^*dose* was observed for markers in all allele frequency classes (Fig. 3a) but it was greater (from 0.688 to 0.818; 19%) for low-frequency (MAF < 5%) than for common variants (MAF > 5%; 0.909 to 0.965; 6%).

**Figure 3.**
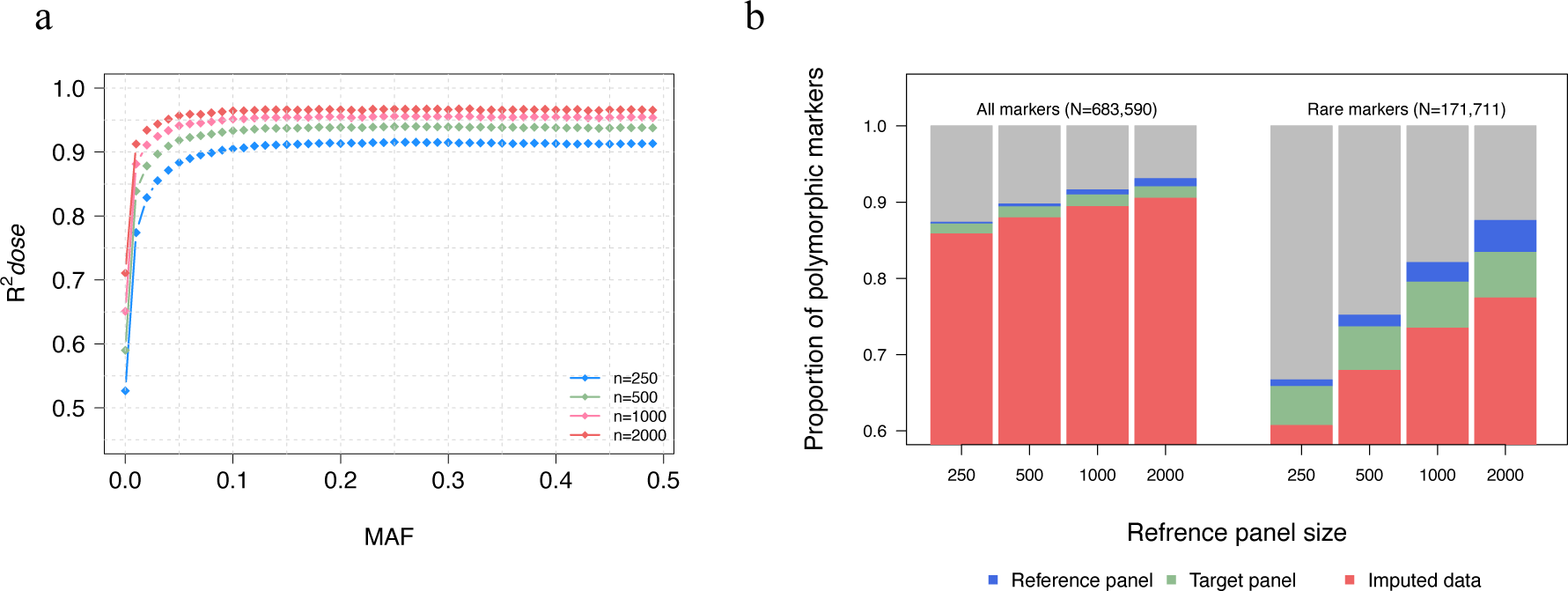
Effect of reference panel size on imputation quality. Imputation accuracy (R2*dose*) for markers grouped in MAF bins when imputing with reference panels of different sizes (a). Proportion polymorphic markers (all and rare; MAF < 2%) in the full data set (n=3938) that are polymorphic in the reference, target and imputed data, when imputing with reference panels of varying sizes (b).

The correlation-based empirical accuracy metric R^*2*^*dose* does not reflect the realized marker density (number of informative markers) in the imputed data set. Therefore, we studied the marker density in the imputed dataset when the genotypes were imputed using different reference panels. Larger reference panels captured a greater fraction of genetic variation in the population, allowing the imputation of a larger number of variants (Figure 3b and Table 2). In the smallest (n=250) and the largest (n=2000) reference panels, 597,095 and 636,201 markers were polymorphic, respectively, of which 595,609 and 629,025 were also polymorphic in the 1938 target animals.

**Table 2.** Number of polymorphic and monomorphic markers in reference, target and imputed data

| Ref<sup>a</sup> | Marker | Number of markers |  |  |  |  |
| --- | --- | --- | --- | --- | --- | --- |
|  |  | Polymorphic in reference panel | Polymorphic in target panel | Monomorphic in target panel | Polymorphic in imputed data | Falsely imputed<sup>b</sup> |
| 250 | All | 597,095 | 595,609 | 1486 | 586,701 | 526 |
|  | Rare<sup>c</sup> | 114,516 | 113,030 | 1486 | 104,222 | 526 |
| 500 | All | 613,649 | 610,987 | 2661 | 601,082 | 711 |
|  | Rare<sup>c</sup> | 129,107 | 126,446 | 2661 | 116,619 | 711 |
| 1000 | All | 626,122 | 621,698 | 4424 | 611,284 | 860 |
|  | Rare<sup>c</sup> | 140,931 | 136,507 | 4424 | 126,155 | 860 |
| 2000 | All | 636,201 | 629,025 | 7176 | 618,660 | 990 |
|  | Rare<sup>c</sup> | 150,407 | 143,231 | 7176 | 132,931 | 990 |

<sup>a</sup>Reference panel size
<sup>b</sup>Number of markers that were imputed as polymorphic but were monomorphic in the target animals (considering the true genotypes)
<sup>c</sup>MAF < 2%

When we reduced the marker density of the target animals to the content of the BovineSNP50 BeadChip genotyping array and imputed the missing genotypes using the respec^−^ tive reference panels, 586,701 and 618,660 markers remained informative. The increase in the imputed marker density with larger reference panels was mainly (∼90%) due to a higher number of low-frequency variants (Table 2). The largest reference panel contained ∼31% more (150,407 vs. 114,516) rare alleles (MAF < 2%) than the smallest reference panel (n=250 animals), of which 132,931 (92.2%) and 104,222 (92.8%) remained informative post-imputation. The additional markers available in the imputed data when genotypes were inferred from the largest reference panels (compared to the smallest reference panel) were imputed with the mean and median R^2^*dose* of 0.674 and 0.740 with 42% of the markers being imputed with R^2^*dose* > 0.80.

A subset of rare (MAF < 2%) markers segregating in the reference panel was monomorphic/fixed in the target animals (considering the true genotypes). Some of them turned out to be polymorphic post-imputation likely due to erroneously imputed genotypes. The absolute number of monomorphic markers in the target panel that were polymorphic post-imputation increased (526 vs. 990 for n=250 and 2000 respectively), but the percentage (35% and 14% respectively) decreased with the size of the reference panel. Interestingly, Minimac3 reported moderately high R*sq* values for these markers, increasing with the size of the reference panel (0.640, 0.683, 0.718, and, 0.740 using reference panels with 250, 500, 1000 and 2000 animals respectively).

### Effect of target panel size on imputation accuracy and markers density

We investigated the effect of target panel size on imputation accuracy and number of polymorphic markers after imputation using a target panel of 500 (base) animals. The 50K genotypes of the base animals were phased together with 0, 600, 1200, 1800, or 2400 additional animals, and subsequently imputed to 777K using the same reference panel of 1000 animals.

The mean squared correlation between true and imputed allele dosages (R^*2*^*dose*) increased when genotypes were imputed into haplotypes that were inferred in larger target panels (Figure 4a and Table 3). For instance, R^*2*^*dose* increased from 0.903 to 0.913 (1.1%) when 2400 additional animals were included for haplotype inference. Increase in accuracy was observed for markers in all allele frequency classes (Fig. 4a). Rare variants (MAF < 2%) benefited the most from large target panels. For common variants, the marginal increase in accuracy was low beyond 600 additional animals (Table 3).

**Table 3.** Imputation accuracy (R^*2*^*dose*) in 500 target samples when additional animals were used for haplotype inference

|  | Number of additional samples used for the phasing of 500 target base animals |  |  |  |  |
| --- | --- | --- | --- | --- | --- |
|  | 0 | 600 | 1200 | 1800 | 2400 |
| **Rare** ($MAF < 2\%$) | 0.748 | 0.760 | 0.765 | 0.767 | 0.768 |
| **Low-frequency** ($MAF < 5\%$) | 0.804 | 0.815 | 0.819 | 0.820 | 0.821 |
| **Common** ($MAF > 5\%$) | 0.948 | 0.952 | 0.953 | 0.954 | 0.954 |
| **Global** | 0.903 | 0.910 | 0.912 | 0.913 | 0.913 |

**Figure 4.**
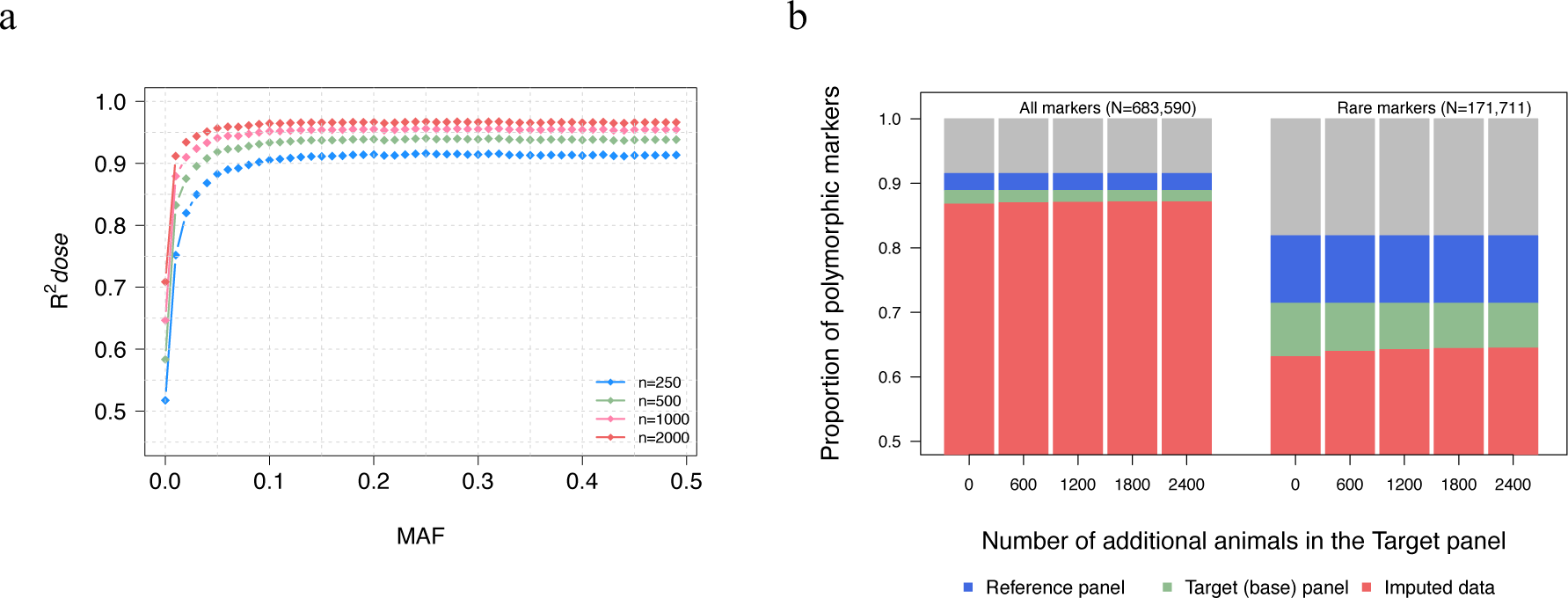
Effect of target panel size on imputation quality. Imputation accuracy (R^*2*^*dose*) for markers grouped in MAF bins when genotypes were imputed into the haplotypes of 500 target samples that were inferred together with additional samples (a). Proportion of polymorphic markers (all and rare; MAF <2%) in the full data (n=3938) that are polymorphic in the reference (n=1000), base target (n=500) and, imputed data (base), when genotypes were imputed into haplotypes of 500 base animals inferred with different number of additional samples (b).

In the 1000 reference animals, 625,538 markers were polymorphic of which 607,560 were also polymorphic in the 500 base animals. When we combined the base animals with different numbers of additional animals (0, 600, 1200, 1800, and 2400) and imputed the missing genotypes, the number of markers that were polymorphic in the imputed data increased with the number of additional animals used. For instance, the number of polymorphic markers in the imputed data increased by 2331 markers (from 593,239 to 595,570) when 2400 additional animals were considered to infer the haplotypes of the 500 base animals. Most (n=2323) of them were rare markers segregating at MAF < 2% (Fig. 4b).

Of 17,978 rare markers that had MAF less than 2% in the reference panel, 14.6% were monomorphic in the true genotypes of the target panel. However, some of these rare markers were polymorphic in the imputed dataset likely due to erroneously imputed genotypes. The fraction of fixed markers in the target data that turned polymorphic after imputation increased with the number of additional animals used for phase inference in target animals (9.5 and 10.5% using 0 and 2400 additional animals for haplotype inference, respectively) (Table 4).

**Table 4.** Number of polymorphic and monomorphic markers in n=1000 reference and n=500 target animals when different numbers of additional animals were included for phase inference in the 500 target animals

| Add<sup>a</sup> | Marker | Number of markers |  |  |  |  |
| --- | --- | --- | --- | --- | --- | --- |
|  |  | Reference panel | Polymorphic in target panel | Monomorphic in target panel | Polymorphic in imputed target panel | Falsely imputed<sup>b</sup> |
| 0 | All | 625,538 | 607,560 | 17,978 | 593,239 | 1702 |
|  | Rare<sup>c</sup> | 140,584 | 122,606 | 17,978 | 108,379 | 1702 |
| 600 | All | 625,538 | 607,560 | 17,978 | 594,682 | 1831 |
|  | Rare<sup>c</sup> | 140,584 | 122,606 | 17,978 | 109,818 | 1831 |
| 1200 | All | 625,538 | 607,560 | 17,978 | 595,150 | 1856 |
|  | Rare<sup>c</sup> | 140,584 | 122,606 | 17,978 | 110,282 | 1856 |
| 1800 | All | 625,538 | 607,560 | 17,978 | 595,433 | 1880 |
|  | Rare<sup>c</sup> | 140,584 | 122,606 | 17,978 | 110,568 | 1880 |
| 2400 | All | 625,538 | 607,560 | 17,978 | 595,570 | 1886 |
|  | Rare<sup>c</sup> | 140,584 | 122,606 | 17,978 | 110,702 | 1886 |

<sup>a</sup> Number of additional animals
<sup>b</sup>Number of markers that were imputed as polymorphic but were monomorphic in the base target (considering the true genotypes).
<sup>c</sup>MAF < 2%

### Reliability of the model-based quality score from Minimac3

Next, we compared the model-based quality score reported by Minimac3 (R*sq*) to our three empirical accuracy estimates -- (i) squared Pearson’s correlation coefficient between true genotypes and imputed allele dosages (R^2^*dose*) (ii) squared Pearson’s correlation coefficient between true and imputed “best-guess” genotypes (R^2^*gt*), and (iii) proportion of correctly imputed genotypes (concordance rate – CR). Irrespective of the size of the reference panel and MAF of the imputed variants, the R*sq* values were higher than R^2^*dose* and R^2^*gt* but lower than CR values (Table 1, Additional file 4). However, the discrepancy between these metrics decreased when either the MAF of the imputed variants or the size of the reference panels increased (Additional file 5, Table 1).

The R*sq* values reported by Minimac3 were highly correlated with R^*2*^*dose* (0.77) and R^*2*^*gt* (0.75) but not with CR (0.03) when a reference panel of 2000 samples was used to impute 1939 target samples. The very low correlation between R*sq* and CR was mainly due to large discrepancies of both metrics for markers with low MAF. The mean correlations of R*sq* with R^*2*^*dose*, R^*2*^*gt*, and CR (excluding markers with MAF < 0.5%) increased from 0.80, 0.81 and, 0.33 to 0.84, 0.84 and, 0.61, respectively, when the reference panel size increased from 250 to 2000 (Fig. 5).

**Figure 5.**
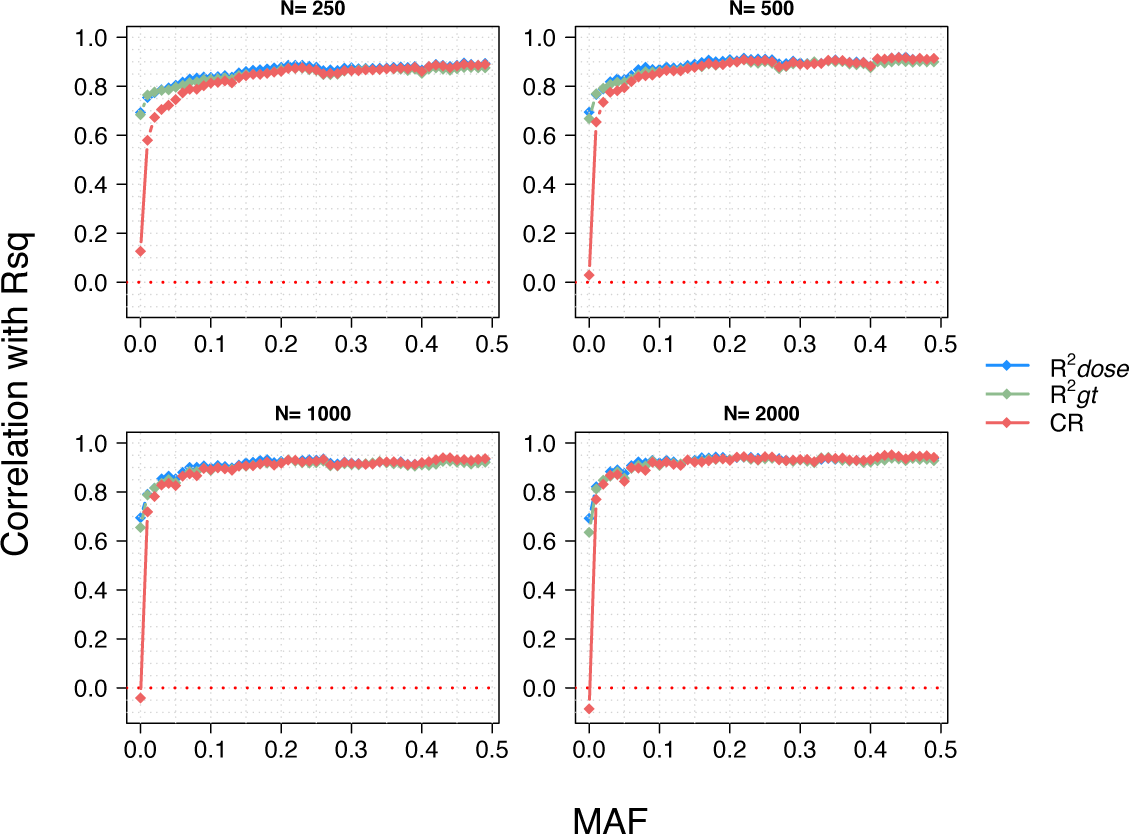
Correlation between model-based imputation quality score from Minimac3 (R*sq*) and empirical measures of imputation accuracy. Correlation of three empirical measures of imputation accuracy (R^*2*^*dose*, R^*2*^*gt*, CR) with the model-based imputation quality score (R*sq*) of Minimac.

These correlations were also stronger for common than for the rare variants; when imputing with the largest (n=2000) reference panel, the correlations of R*sq* with R^*2*^*dose*, R^*2*^*gt* and CR were 0.81, 0.81 and, 0.64 and 0.93, 0.92 and, 0.84 for low-frequency (MAF < 5%) and common (MAF > 5%) variants, respectively.

## Discussion

Phasing and imputation accuracies of high-density genotypes using the ARS-UCD1.2 bovine reference genome assembly have been investigated in different *Bos taurus taurus* breeds (11). In *Bos taurus indicus* cattle, the accuracy of imputing 50K genotypes to higher density has been investigated only using the UMD3.1 assembly of the bovine genome (23, 24). To the best of our knowledge, our study is the first to evaluate the accuracy of imputing 50K to 777K genotypes in a *Bos taurus indicus* cattle breed using the ARS-UCD1.2 assembly of the bovine genome (21).

Higher contiguity of the ARS-UCD1.2 genome assembly might improve haplotype inference in regions that contained phasing and imputation errors in the previous assembly (UMD3.1) (7,10,18). While the global accuracy of imputation was identical for both assemblies, we detected a number of segments for which the accuracy of imputation differed greatly. Moreover, we identified fewer putatively misplaced SNPs on ARS-UCD1.2 compared to UMD3.1 and observed a substantial improvement in imputation accuracy for 60% of misplaced SNPs on UMD3.1 when they were mapped to the new build. However, we also identified SNPs that had considerably higher imputation accuracy when aligned to UMD3.1 than to ARS-UCD1.2 indicating that some physical ARS-UCD1.2 coordinates of the 777K markers are wrong. A limitation of our study is that we considered only SNPs interrogated with commercial genotyping chips. These SNPs suffer from ascertainment bias as they are predominantly located in more accessible regions of the genome (25). Hence differences in imputation accuracy between the two builds might be higher than observed in our study when imputing whole-genome sequence variant genotypes. A haplotype-resolved Brahman genome assembly (26) might further improve the accuracy of imputing genotypes for animals from indicine breeds, particularly within regions that are not co-linear between *Bos taurus taurus* and *Bos taurus indicus* assemblies.

The accuracy of imputing Illumina BovineHD genotypes was assessed previously in indicine cattle. Accuracy greater than 0.957 was obtained when reference panels of 793 and 171 Nellore and Gyr animals were used to impute 50K genotypes to 777K (23,24). Using map positions of the UMD3.1 assembly and the software FImpute (4), these studies assessed the correlation between true and imputed (best-guess) genotypes (R*gt*). Using a pre-phasing-based approach that does not explicitly consider pedigree information, we obtained a lower mean imputation accuracy (R*gt*) of 0.952 in Nellore cattle, although the number of reference animals (n=2000) was considerably higher and the pre-phasing-based approach applied in our study is expected to be at least as accurate as FImpute (27). The slightly lower accuracy obtained in our study likely resulted from including more low-frequency markers, that are difficult to impute, rather than from differences in the imputation software used. Applying the same MAF filter used in the previous studies (MAF > 2%), we indeed obtained a comparable accuracy (R*gt*) of 0.976 using the ARS-UCD1.2 assembly with pre-phasing-based genotype imputation.

Accurate genotype imputation requires large reference panels that are genetically related to the target animals (3,7,28). In our study, the mean accuracy of imputation (R^*2*^*dose*) increased by 8.52% when the size of the reference panel increased from 250 to 2000. Our findings also show that the size of the reference panel is correlated to the number of informative markers, i.e., polymorphic, in the imputed data set. Large reference panels facilitate imputing genotypes for true rare variants. Importantly, genotypes for additional variants that were only available when the largest reference panel was considered, were imputed with moderately high accuracy (mean R^*2*^*dose*=0.674). Correlation-based accuracy estimates (R^*2*^*-dose* and R^*2*^*gt*) do not reflect the number of monomorphic sites post imputation, thus the overall benefit from large reference panels is even higher than indicated by these measures (12,29)

Our approach allowed for an indirect investigation of the effect of sample size on phasing accuracy by studying the accuracy of imputing genotypes into the inferred haplotypes. We show that the size of the target panel also influences pre-phasing-based imputation quality. Both imputation accuracy and marker density in the target animals increased when their haplotypes were inferred in larger cohorts. Haplotype inference using statistical methods benefits from large cohorts (15) due to better modeling of linkage disequilibrium and increased prevalence of identical by descent tracts (16). Although the pre-phasing-based strategy employed here is computationally efficient (and accurate when using larger reference and target panels), our results show that care needs to be taken when it is applied to impute genotypes into smaller cohorts. The new paradigm of reference-based phasing implemented in Eagle2 and SHAPEIT3 (30), or use of pedigree information (e.g., with FImpute (4) or LINKPHASE3 (31)) might improve phase inference in small reference and target panels. As noted by Loh and coworkers (16), the reference panel used for imputation could also be used as a reference panel for phase inference. If the size of the target panel increases, it seems advisable to infer target haplotypes and impute genotypes in the pooled data again instead of imputing only the new data. Our results may be generalized to other applications of genotype phasing as they highlight that the sample size needs to be considered to choose an appropriate phasing method.

Many genotype imputation software tools estimate a SNP-level model-based quality score to indicate imputation accuracy. This value may be used to detect and discard poorly imputed variants from downstream analyses (28,32). We compared the model-based imputation quality score from Minimac3 (R*sq*) to three empirical measures of imputation accuracy (R^*2*^*dose*, R^*2*^*gt*, and CR). In agreement with previous studies that investigated the model-based accuracies from Beagle (33) and Minimac3 (22), we found that R*sq* values were highly correlated with correlation-based empirical measures (R^*2*^*dose*, R^*2*^*gt)*. The correlation between empirical and model-based quality scores increased with the size of the reference panel. However contrary to the results from a previous study (22), we found R*sq* values to be systematically higher than the R^*2*^*dose* and R^*2*^*gt* for makers across all allele frequency classes. The third empirical measure, CR was not correlated with R*sq* when all markers were considered. This measure, unlike the first two, does not account for the frequency of imputed alleles and considerably overestimates the accuracy of imputation for rare variants (34,35). However, the correlation between CR and model-based quality score is also high for variants with MAF greater than 5%.

The calculation of empirical accuracy using cross-validation is computationally intensive. Our results show that the model-based quality score is highly correlated with empirical correlation-based measures. However, Minimac3 reported moderately high R*sq* values (increasing with the size of the reference panel used; > 0.640) for variants that were falsely imputed (i.e. monomorphic in the target panel, considering the true genotypes, but polymorphic in the imputed data). Because thresholds between 0.3 and 0.6 are commonly used to discard markers of insufficient imputation quality (36) these erroneously imputed variants would qualify for downstream analyses. Variants that are imputed into target animals although they are actually not polymorphic, may result in conflicting results in association studies (37). Our findings show that model-based imputation quality scores, although useful, need to be treated with caution, particularly for low-frequency and rare variants that have been imputed from less informative reference panels.

## Conclusions

The improvements in the new assembly of the bovine genome (ARS-UCD1.2) affected the imputation of only a small fraction of SNPs present on the BovineHD SNP chip. The global accuracy of imputation was virtually identical to the previous build (UMD3.1). Accuracy of imputation and the number of informative markers in the imputed data benefit from large reference panels. The size of the target panel also has an influence on both metrics when genotypes are inferred using pre-phasing-based approaches. The model-based imputation quality scores from Minimac3 (R*sq*) were correlated with the empirical correlation-based estimates of accuracy. These values can be used to filter out poorly imputed variants but their reliability is low for rare and low-frequency variants particularly when genotypes are imputed from small reference panels.

## Methods

### Genotype data

We considered microarray-derived SNP genotypes of 4098 Nellore (*Bos taurus indicus)* cattle from farms participating in the DeltaGen^®^ and Paint^®^ (CRV Lagoa) breeding programs in Brazil. All animals were genotyped using the high-density Illumina BovineHD BeadChip (Illumina, Inc., San Diego, CA, USA), comprising 777,962 SNPs. The map positions of the SNPs were determined according to the *Bos taurus taurus* reference genome assembly UMD3.1 (38). Quality control (QC) on the raw genotype data was performed using the PLINK (v1.9) software (39). Animals and SNPs with a call rate < 95% were not considered. SNPs that were annotated to non-autosomal chromosomes or SNPs for which the map positions were unknown were discarded as well as SNPs that deviated from Hardy-Weinberg proportions (*p* < 10^−5^). After filtering, 3938 animals with genotypes for 684,561 autosomal SNPs remained in the dataset. Sporadically missing genotypes (max 5%) were imputed using Beagle5 (40)

### Liftover to ARS-UCD1.2

We used the ARS-UCD1.2 (liftover) coordinates (available from https://www.animalgenome.org/repository/cattle/UMC_bovine_coordinates/) to determine the physical coordinates of the Illumina BovineHD BeadChip markers according to the ARS-UCD1.2 genome assembly. Physical coordinates of 774,091 SNPs were available for the ARS-UCD1.2 assembly. Using the QC parameters as detailed above, we excluded 90,501 SNPs and 160 individuals. The final dataset (ARS-UCD1.2) included 3938 animals with genotypes at 683,590 autosomal SNPs.

### Imputation procedure and assessment of accuracy

To assess the accuracy of imputation for different compositions of reference and target panels and two bovine genome assemblies (see below), we carried out a ten-fold cross-validation using the 3938 animals. For each fold, we randomly grouped animals into target and reference panels. The marker density in the target panels was reduced to match the BovineSNP50 (version 2) BeadChip comprising 56,206 SNPs. Subsequently, the genotypes of the target panels were imputed to higher density using information from the reference panel using a pre-phasing-based imputation workflow. The Eagle v2.4 (41) software was used to infer haplotypes for the reference and target panels and Minimac3 (13) was used to impute genotypes.

Minimac3 provides imputed allele dosages (continuously distributed values between 0 and 2) and best-guess genotypes (discrete values of 0, 1, or 2). We assessed empirical imputation accuracy as the squared Pearson’s correlation coefficient between either true genotypes and imputed allele dosages (R^*2*^*dose*), or true genotypes and best-guess genotypes (R^*2*^*gt*). We also calculated the proportion of correctly imputed genotypes (concordance rate – CR). R^*2*^*-dose* values were used to assess imputation quality in different imputation scenarios and all three empirical accuracies were used to investigate the reliability of the model-based quality score estimate from Minimac3 (R*sq*).

Values for R^*2*^*dose* and R^*2*^*gt* cannot be calculated for markers that are imputed to the major allele in all target animals because the imputed doses and “best-guess” genotypes for such markers have zero variance. Thus, the loss of such otherwise segregating markers (usually at very low MAF) post-imputation is not taken into account by the correlation-based imputation accuracy measures. To study the effect of reference and target panel composition on the realized marker density in the imputed dataset, we also calculated the proportion of markers that are imputed to the major allele in all target samples but were segregating in the real data set.

### Identification of misplaced SNPs

We analyzed the accuracy of genotypes imputed from the largest reference panel to identify regions with poor imputation quality -- due most likely to misplacement of SNPs -- when genotypes were aligned to the two bovine genome assemblies. We partitioned autosomes into segments of 100 kb and counted the number of poorly imputed (MAF > 1% and R^2^*dose* < 0.8) SNPs. All SNPs in segments with more than three poorly imputed SNPs were considered as putatively misplaced.

### Test Scenarios

#### Effect of reference and target panel

We used 3938 animals with genotypes for 683,590 markers aligned to the ARS-UCD1.2 assembly to investigate the effect of reference and target panel sizes on imputation quality. To study the effect of the reference panel, we imputed a target panel of 1938 animals from 50K to 777K using reference panels with 250, 500, 1000, and 2000 randomly sampled animals in 10 replicates. To study the effect of target panel size, we imputed genotypes in pre-phased target panels of varying sizes using a reference panel of fixed size (n=1000). Genotypes for 500 animals (base panel) were phased with an additional of 0, 600, 1200, 1800 or 2400 randomly sampled animals. The resulting pre-phased target panels respectively containing 500, 1100, 1700, 2300, and 2900 animals were imputed to higher density using a phased reference panel of 1000 animals. The imputation was carried out in 10-fold cross-validation, and the accuracy of imputation was assessed only for the 500 animals in the base panel.

#### Comparison between UMD3.1 and ARS-UCD1.2 genome assemblies

We compared the accuracy of imputation in SNP genotypes aligned either the UMD3.1 or ARS-UCD1.2 genome assembly by imputing 1938 animals from 50K to 777K using a reference panel of 2000 animals in ten-fold cross-validation (see above). Target and reference animals were selected randomly. The global imputation accuracies were reported as mean R^*2*^*dose* (see above) from all autosomal markers across all replicates. We also studied the difference in accuracies for

i. “newly mapped” markers i.e., markers that are unmapped when aligned to UMD3.1 and on autosomes when aligned to ARS-UCD1.2,
ii. markers aligned to different chromosomes for the assemblies, and
iii. markers identified as putatively misplaced.

## Availability of data and materials

The ARS-UCD1.2 (liftover) coordinates are available from https://www.animalgenome.org/repository/cattle/UMC_bovine_coordinates/. Our research was accomplished using data from commercial farms and Nellore breeding programs that are not publicly available.

## Acknowledgements

This study was financed in part by the Coordenação de Aperfeiçoamento de Pessoal de Nível Superior – Brasil (CAPES) – Finance Code 001.

## Author information

### Authors contributions

NKK and HP conceived and designed the study. ICH and NKK performed the analyses. RBC and LGA provided genotypes, read and revised the manuscript. ICH, NKK and HP wrote the manuscript. All authors have agreed on the content of the manuscript.

### Corresponding author

Correspondence to Naveen Kumar Kadri

## Ethics declarations

### Ethics approval and consent to participate

Not applicable.

### Consent for publication

Not applicable.

