## Additional file 1 for "Investigating the accuracy of imputed genotypes in Nellore cattle using the ARS-UCD1.2 assembly of the bovine genome"

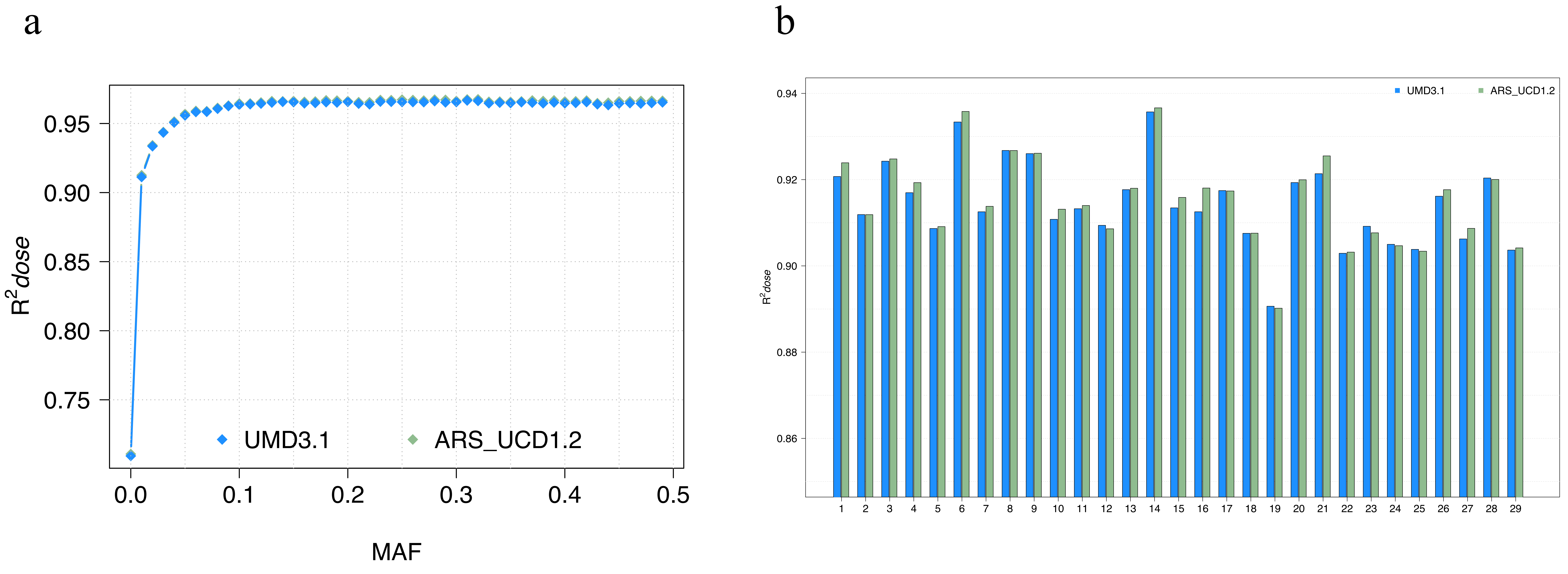


**Additional File 1. Imputation accuracy according to reference genome assembly.**
Accuracy of imputation for markers grouped in MAF bins when aligned to the two Bovine reference assemblies (ARS-UCD1.2 and UMD3.1) (a). The mean imputation accuracy for markers on the 29 autosomes when aligned to the two Bovine assemblies (b).
