## Additional file 4 for "Investigating the accuracy of imputed genotypes in Nellore cattle using the ARS-UCD1.2 assembly of the bovine genome"

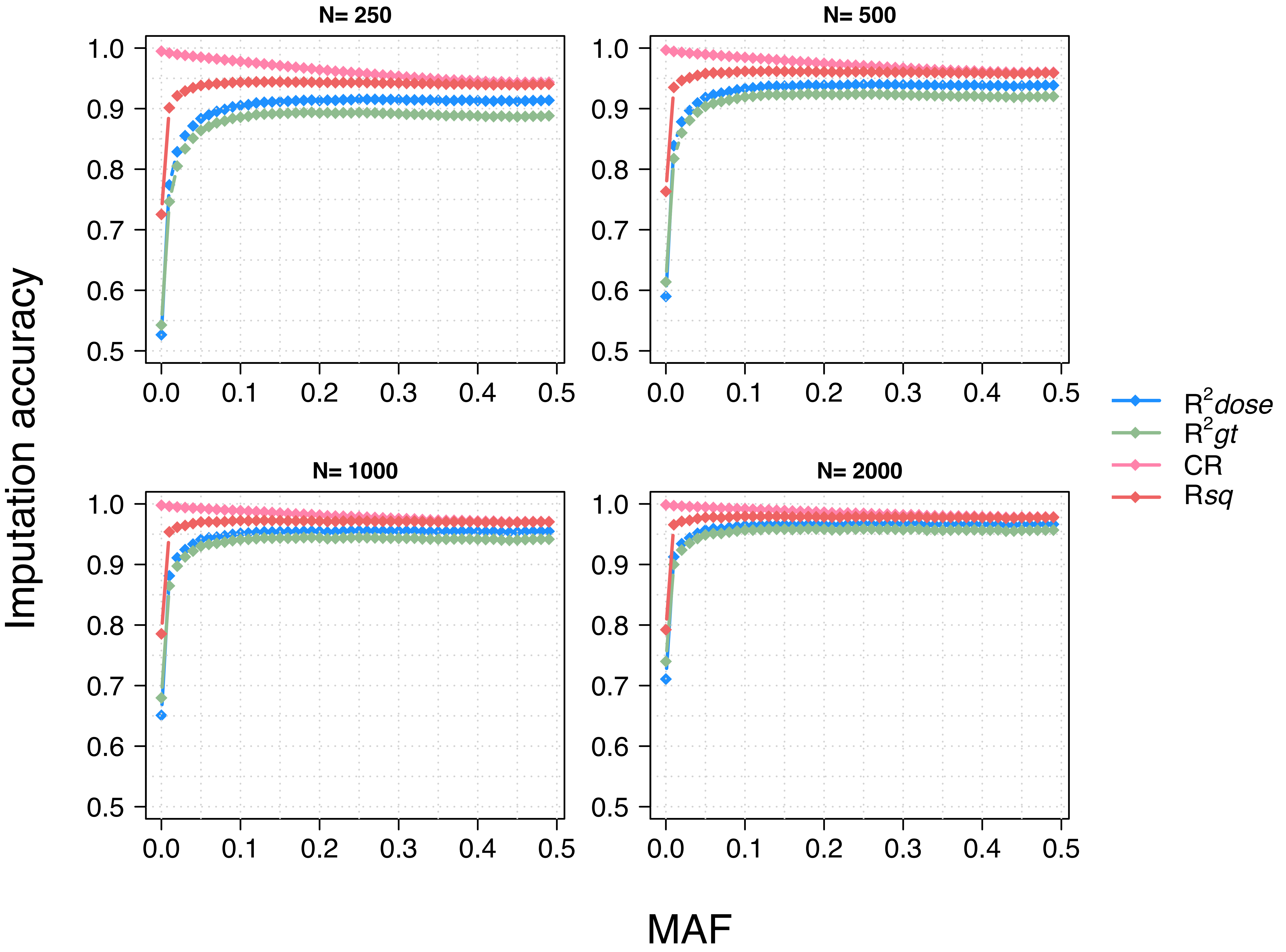


**Additional File 4. Model-based imputation quality score from Minimac3 (R*sq*) and empirical measures of imputation accuracy.**
Empirical measures of accuracy and the model-based imputation quality scores from Minimac3 (R*sq*) for markers grouped in MAF bins when imputing with reference panels of varying sizes.
