## Additional file 5 for "Investigating the accuracy of imputed genotypes in Nellore cattle using the ARS-UCD1.2 assembly of the bovine genome"

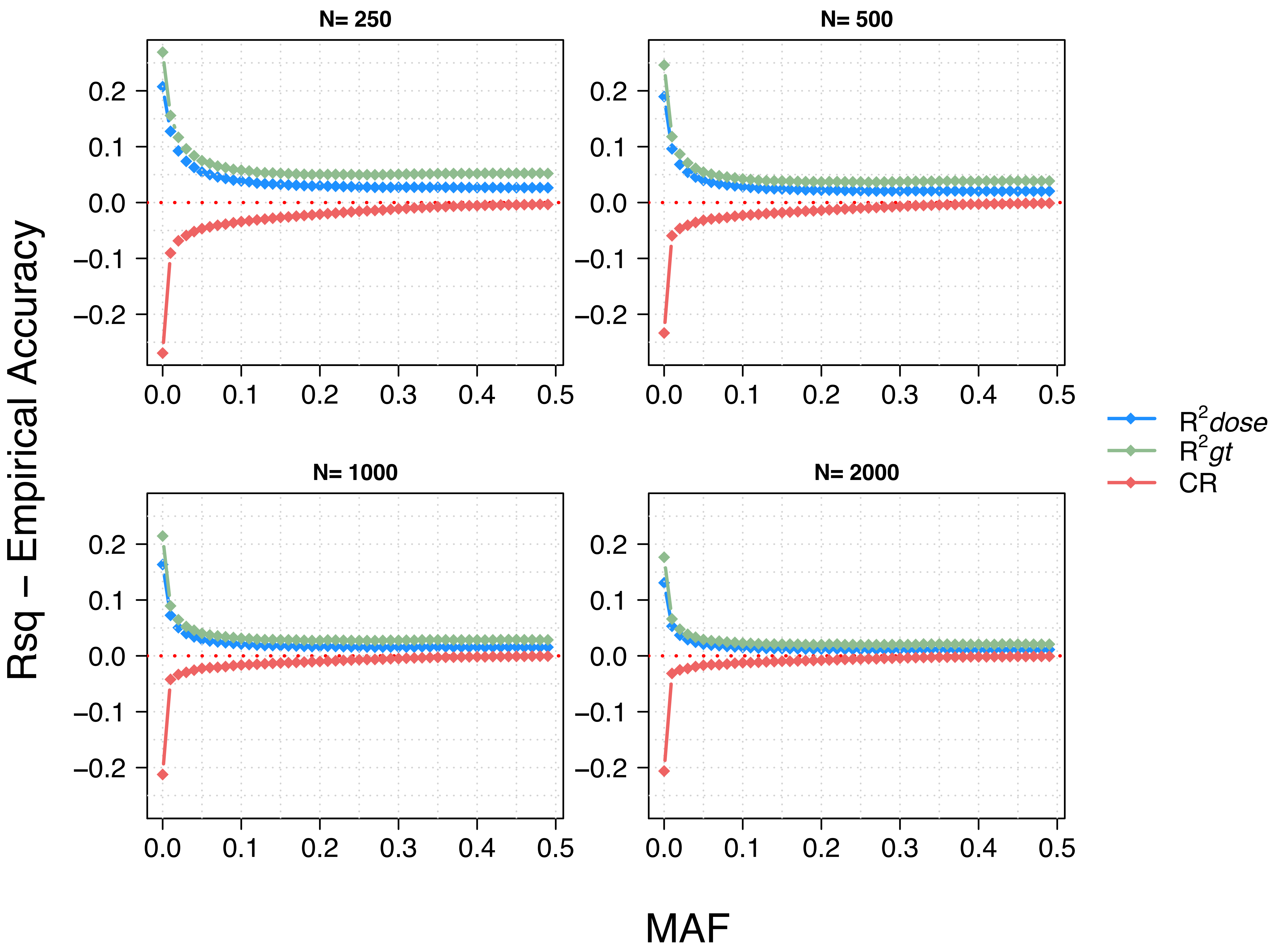


**Additional File 5. Discrepancy between the model-based imputation quality scores from Minimac3 (R*sq*) and empirical measures of accuracy.**

Discrepancy between the model-based imputation quality scores from Minimac3 (R*sq*) and empirical measures of accuracy for markers grouped in MAF bins when imputing with reference panels of varying sizes.
